# In Utero Hematopoietic Stem Cell Transplant for Fanconi Anemia

**DOI:** 10.1101/2024.05.09.592452

**Authors:** Leah Swartzrock, Carla Dib, Morgane Denis, Hana Willner, Katie Ho, Ethan Haslett, Mark R. Krampf, Anna Girsen, Yair J. Blumenfeld, Yasser Y. El-Sayed, Maria G Roncarolo, Tippi C. MacKenzie, Agnieszka D. Czechowicz

## Abstract

Fanconi Anemia (FA) is an inherited DNA-repair deficiency caused by mutations in diverse *Fanc* genes that leads to bone marrow failure and malignancies. FA disease begins at early embryonic stages, and while FA prenatal testing has long been available, no fetal therapies for FA currently exist. Postnatally, FA hematologic disease can be cured through allogeneic hematopoietic stem cell transplantation (HSCT); however, this requires chemotherapy and/or irradiation-based conditioning which amongst various side-effects also increases likelihood of malignancies later in life in these fragile patients. Given fetal immune tolerance and the competitive advantage of healthy hematopoietic stem and progenitor cells (HSPCs) over failing FA HSPCs, in utero HSCT without conditioning may be an alternative approach to stabilization of the hematopoietic system without conventional toxicities. We performed in utero HSCT using HSPCs from wildtype (WT) donors into two FA mouse models (*Fancd2*^−/−^, *Fanca*^−/−^) and observed robust multi-lineage hematopoietic donor engraftment in homozygous FA mice compared to both heterozygous FA and WT littermates. Upon serial assessments, we also observed increasing donor chimerism up to 94.1%, showcasing the competitive advantage of WT donor HSPCs over FA HSPCs. Given that 1% donor chimerism is predicted to stabilize FA BM, in utero HSCT may be a safe and curative prenatal treatment for all subtypes of FA.

## To the Editor

Fanconi Anemia (FA) is an inherited DNA repair disorder resulting in the inability to repair interstrand crosslinks (ICL)^1,2^. FA is caused by mutations in one of 23 different Fanconi anemia complementation (FANC) genes, with the most frequent mutations occurring in FANCA (∼65%), FANCC (∼10%), and FANCG (∼10%) genes^3^. The enhanced chromosomal fragility of FA individuals leads to progressive bone marrow failure (BMF) and a systemic predisposition to malignancies. Most FA patients develop BMF between 5 and 15 years of age with ∼80% of patients developing BMF by the age of 10^1,3^. Additionally, patients are at very high risk of developing acute myeloid leukemia (AML) and myelodysplastic syndrome (MDS) throughout their lives^1,4^. The only proven curative way to treat the hematologic disease in FA is with allogeneic hematopoietic stem cell transplantation (HSCT) which is required for FA patients with BMF, myelodysplastic syndrome, or acute myeloid leukemia. However, the efficacy of HSCT is often complicated by availability of HLA-matched donors, graft vs host disease (GVHD) or graft rejection. Further, pre-HSCT conditioning poses significant comorbidity in FA patients as it causes direct tissue damage leading to organ damage and infertility, predisposes patients to deadly infections and significantly increases likelihood of malignancies later in life in these hypersensitive patients^4,5^. Early intervention via transplantation of allogeneic HSCs into a FA gestational fetus may be an avenue to overcome limitations of post-natal HSCT and provide hematopoietic stabilization even before the onset of any hematologic symptoms.

Advancements in prenatal diagnostic tools have provided means for early diagnosis of many inherited and congenital diseases, including genetic hematologic diseases. Suspicion for FA in a developing fetus often arises based upon family history and/or detection of congenital abnormalities on ultrasound^6^. Subsequently, FA can be verified by genetic sequencing and/or chromosomal breakage testing on fetal cells obtained by amniocentesis or chorionic villus sampling. Diagnostic tests can be performed as early as 10 weeks’ gestation^6^, which is well within the window of optimal timing for IUHSCT^9^. In utero hematopoietic stem cell transplantation (IUHSCT) has been pioneered as an alternate approach to post-natal HSCT for immunodeficiencies and hemoglobinopathies with a goal to achieve mixed hematopoietic chimerism and donor-specific tolerance without genotoxic myeloablation or immunosuppression^8,9^. This is performed by in utero transplantation of maternally-derived bone marrow cells into the umbilical vein via ultrasound guidance. Despite high technical success and safety of IUHSCT, clinical application has been challenged due limited engraftment which is below therapeutic levels in most diseases that it has been used to date. This is thought to be likely due to competition of host HSCs which cannot be overcome without the use of genotoxic myeloablative conditioning^10^. Excitingly in FA, we have found that only minimal initial donor cell engraftment is necessary to stabilize the failing bone marrow environment due to the competitive advantage of healthy cells over failing FA cells^14^. Given this, we hypothesized that IUHSCT may lead to stabilization of the FA hematopoietic system and provide an early treatment option to FA patients without conventional toxicities.

To test our hypothesis, IUHSCT was first performed in Fanconi Complementation Group D2 (*Fancd2*, C57BL/6N, CD45.2) mice^7,12^. Parental FA carriers (*Fancd2*^*+/−*^) were bred to generate timed FA fetuses. At the E13.5-14.5 stage, fetuses were transplanted via intrahepatic injection with 1×10^6^ CD117+ enriched HSCs from the bone marrow of wild type (WT, C57BL/6J-Ptprc^em6Lutzy^/N, CD45.1) donors as previously detailed^11^ (Figure 1A, S1). Peripheral blood was collected via retro-orbital vein puncture and donor chimerism was assessed by flow cytometry at periodic timepoints. Recipient *Fancd2* mice (CD45.2) and donor WT cells (CD45.1) were differentiated by staining with directly conjugated anti-mouse CD45.2 and CD45.1 antibodies (see Supplemental Materials). At 4-weeks post-IUHSCT, *Fancd2*^−/−^ mice had significantly higher donor engraftment compared to heterozygous *Fancd2*^*+/−*^ (p=0.0007) and WT (p=.0016) littermates (Figure 1B, S2). The average donor granulocyte chimerism was 26% in *Fancd2*^−/−^ mice (n=2) with only 3.5% and 3.1% in heterozygous *Fancd2*^*+/−*^ (n=5) and WT (n=2) littermates respectively. Upon serial assessments, we observed increasing average donor chimerism, showcasing the competitive advantage of WT cells over FA cells. At 30-weeks post-IUHSCT, the average donor granulocyte chimerism increased to 94% in *Fancd2*^-/-^ (p<0.0001), whereas heterozygous *Fancd2*^*+/−*^ and WT littermates had insignificant changes. Multi-lineage assessment showcased parallel increases in donor chimerism over time (Figure 1C) with significant donor immune cell engraftment in both B-cell and T-cell compartments. To assess true HSC engraftment, bone marrow was collected via aspirates at 12 and 28 weeks post-IUHSCT. The long-term HSC (LT-HSC) chimerism reflected similar robust and statistically significant engraftment in *Fancd2*^−/−^ mice (p<0.0001) that increased with time (Figure 1D, S3). Such high donor engraftment of WT cells across hematopoietic lineages has been shown to sufficient for FA disease stabilization and resistance to DNA damage^15^.

**Figure 1:**
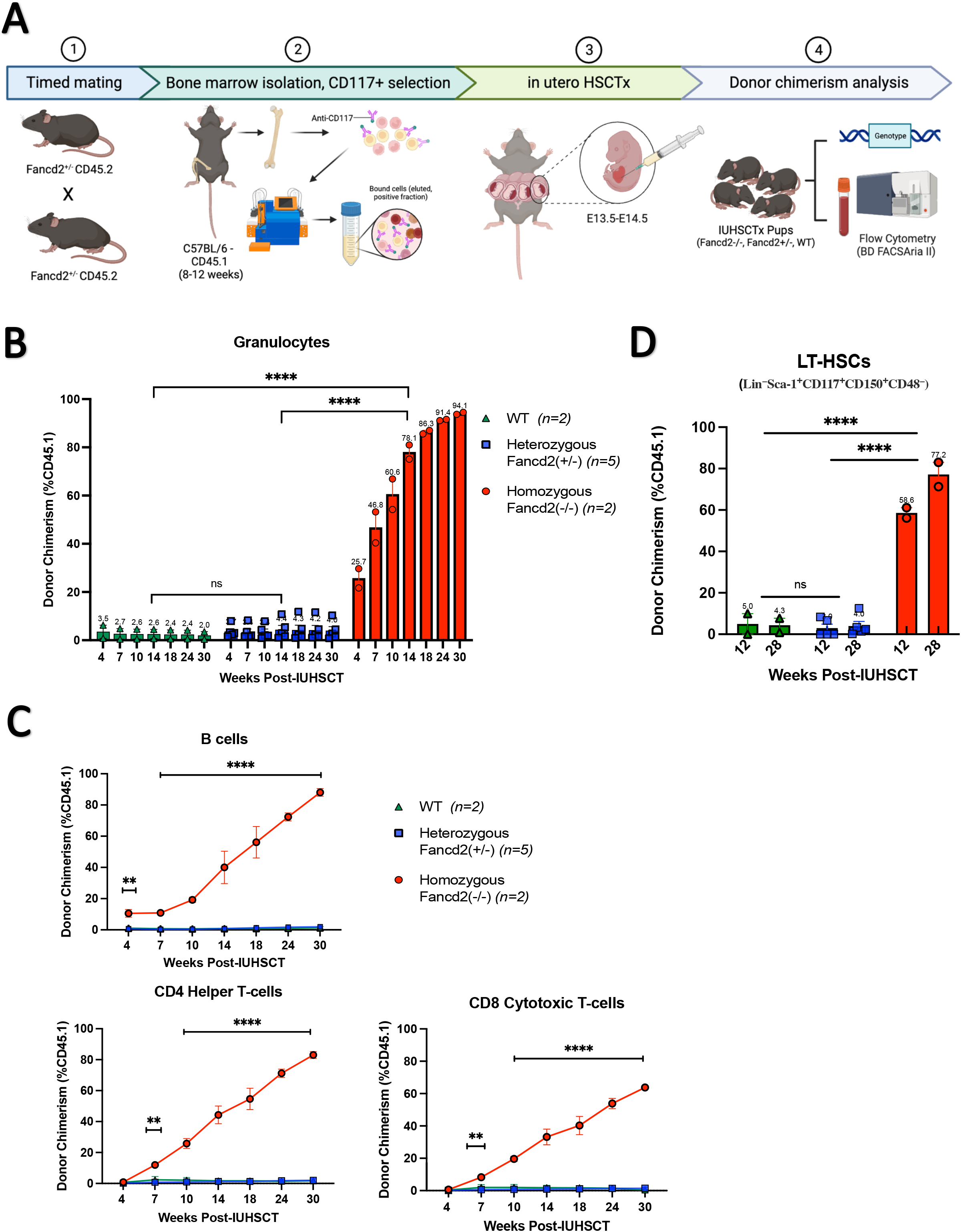
IUHSCT allows for robust donor engraftment in homozygous *Fancd2* ^-/-^ mice with WT competitive advantage over time. (A) Generation of *Fancd2* embryos and in utero transplantation schema. Parental FA carriers (*Fancd2*^*+/*-^ CD45.2) were mated to generate time-dated FA fetuses. Bone marrow from syngeneic wildtype (WT CD45.1) mice was isolated and CD117+ enriched by magnetic separation. At E13.5-14.5, fetuses were transplanted via intrahepatic injection with 1×10^6^ CD117^+^ selected HSCs. Pups were genotyped and donor chimerism was analyzed by flow cytometry (% CD45.1). Peripheral blood and bone marrow chimerism were analyzed at various timepoints post-IUHSCT. (B) Donor granulocyte chimerism, (C) multi-lineage chimerism in B-cells, CD4 Helper T-cells and CD8 Cytotoxic T-cells, and (D) long-term HSC (LT-HSC) chimerism was determined. Data represent mean ± SEM. Comparisons were performed using ANOVA with Tukey’s multiple comparison test and a P-value < 0.05 was considered significant. ****p<0.0001; ***p<0.001; **p<0.01 *p<0.05 ns, nonsignificant.

To explore if similar findings could be observed across diverse FA complementation groups, we further conducted IUHSCT in Fanconi Complementation Group A (*Fanca*, C57BL/6J, CD45.2) mice which represent the most common FA type. FA fetuses were generated, transplanted, and analyzed as previously stated. Similarly, *Fanca*^−/−^ mice had more robust donor engraftment compared to heterozygous *Fanca*^*+/−*^ (p=0.022) and WT (p=0.017) littermates with comparable increased donor granulocyte chimerism (Figure 2A) and multi-lineage chimerism (Figure 2B) in homozygous *Fanca*^−/−^. Bone marrow was collected via aspirate at 25 weeks post-IUHSCT and analyzed for LT-HSC chimerism which was similarly significantly increased in *Fanca*^−/−^ mice (p=0.0157) (Figure 2C). While high engraftment was observed in both *Fanca*^−/−^ and *Fancd2*^−/−^ mice, the increased engraftment in the *Fancd2* model is likely due to the more severe hematopoietic phenotype^12,13^, which consequently results in a greater competitive advantage of the WT donor cells.

**Figure 2:**
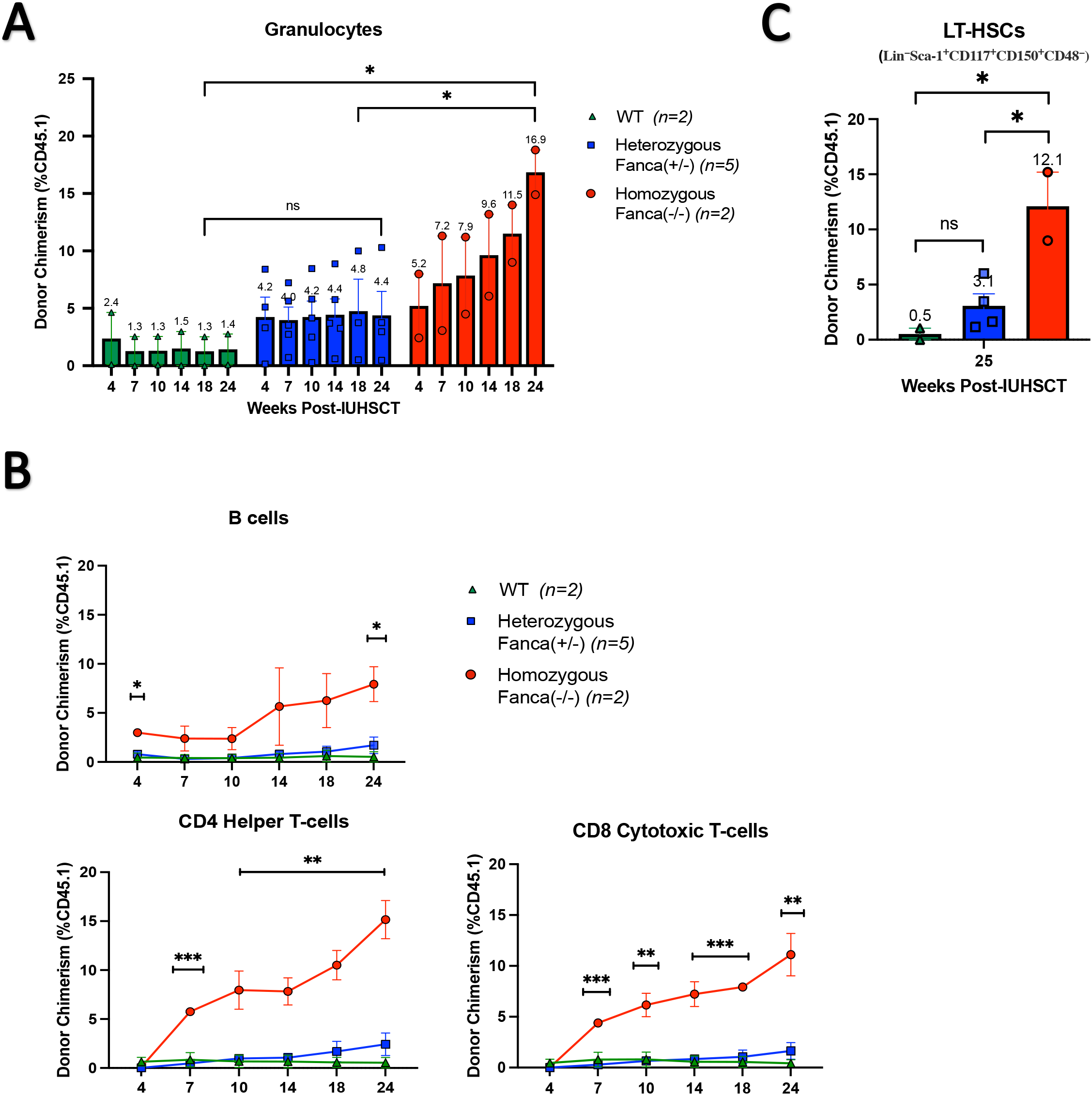
IUHSCT similarly allows for robust donor engraftment in homozygous *Fanca*^-/-^ mice with WT competitive advantage over time. *Fanca* embryos were generated as previously described. At E13.5-14.5, fetuses were transplanted via intrahepatic injection with 1×10^6^ CD117^+^ selected HSCs. Pups were genotyped and donor chimerism was analyzed by flow cytometry (% CD45.1). Peripheral blood and bone marrow chimerism were analyzed at various timepoints post-IUHSCT. (A) Donor granulocyte chimerism, (B) multi-lineage chimerism in B-cells, CD4 Helper T-cells and CD8 Cytotoxic T-cells, and (C) long-term HSC (LT-HSC) chimerism was determined. Data represent mean ± SEM. Comparisons were performed using ANOVA with Tukey’s multiple comparison test and a P value < 0.05 was considered significant. ****p<0.0001; ***p<0.001; **p<0.01 *p<0.05; ns, nonsignificant.

Given prior observations that somatic reversion which likely begins in a single HSC can stabilize the bone marrow in FA patients^15^, our results in two separate FA mouse models with impressive 20-95% donor engraftment through unconditioned IUHSCT highlight that this could become a safe and curative prenatal treatment for all subtypes of FA. The competitive advantage of WT donor HSCs over FA fetal HSCs, combined with the immature and tolerant fetal immune system, allowed for robust engraftment post-IUHSCT in our FA mouse models without the use of genotoxic conditioning or immune suppression. Modern advancements in fetal medicine including high-resolution ultrasound screening and prenatal genetic testing, have improved patient diagnosis of congenital diseases including FA. Our initial findings suggest IUHSCT could be a one-time procedure performed prior to birth that utilizes the tolerant fetal environment to facilitate widespread donor engraftment without any conditioning. Such an approach could be a preventative treatment for the hematologic manifestations of FA, which could subsequently avert the need for other hematologic treatments such as post-natal HSCT which currently has troubling short-term and long-term side effects. IUHSCT is likely to provide a curative treatment option to FA fetuses diagnosed in utero, and further development of such therapies will also additionally enhance FA screening efforts and bring greater awareness of FA amongst the scientific and clinical community.

## Acknowledgments

The authors thank Prof. Kenneth I. Weinberg, Stanford University for his generous gift of the *Fancd2*^*−/−*^ mice, which originated from Prof. Marcus Grompe at Oregon Health and Science University. The authors thank Prof. Irving L. Weissman at Stanford University and Dr. Jinyi Xiang within his laboratory for their collaboration to generate the *Fanca*^−/−^ mice. The authors also thank Prof. David Scadden at Massachusetts General Hospital/Harvard University for his generous gift of the donor C57BL/6N-CD45.1^STEM^ mice. They also thank Amelia Scheck and Cynthia Klein at Stanford University for their help with laboratory management and the Stanford University Stem Cell Institute FACS Core for flow cytometry access and assistance. Additionally, the authors thank the Stanford University School of Medicine Dunlevie Maternal-Fetal Medicine Center for Discovery, Innovation and Clinical Impact and the laboratory of Prof. Tippi MacKenzie at UCSF including Dr. Marisa Schwab and Maria Clarke within her laboratory for their collaboration and guidance of this project. Figures were created with BioRender and GraphPad.

## Authors’ contributions

L.S., A.G., Y.J.B., Y.Y.E., M.G.R., T.C.M., and A.D.C. conceptualized and designed the studies; L.S., H.W., M.D., C.D., and M.R.K. performed the research, collected results, and analyzed data; E.H. and K.H. managed animal colonies and provided technical assistance; L.S., M.G.R., T.C.M., and A.D.C. interpreted data; and L.S. and A.D.C. wrote the manuscript. A.D.C. provided mentorship to the first author.

## Data-sharing statement

Data that support the findings of this study are available from the corresponding author upon reasonable request.

