## Supplemental for "In Utero Hematopoietic Stem Cell Transplant for Fanconi Anemia"

**Supplemental Figures:**

**S1: IUHSCT graft optimization through comparison of different HSC enrichment strageties in wildtype (WT) mice.** Bone Marrow from syngeneic wild type (WT CD45.1) mice was isolated and Lineage depleted or CD117+ enriched by magnetic separation. At E13.5-14.5, WT fetuses were transplanted via intrahepatic injection with either $1x10^6$ Lineage$^-$ or $0.5x10^6$ CD117$^+$ selected HSCs. Peripheral blood was analyzed at various timepoints and bone marrow chimerism was analyzed at 16 weeks post-IUHSCT. (A) Donor granulocyte chimerism and (B) long-term HSC (LT-HSC) chimerism was determined.

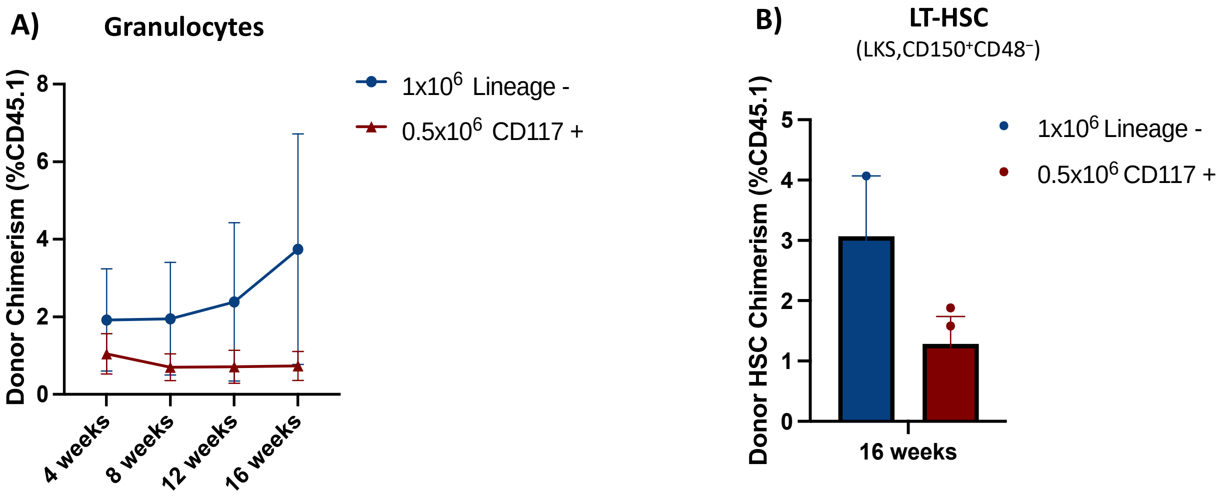


**S2: Representative gating strategy for peripherial blood chimerism.** Post-IUHSCT, donor engraftment in the peripheral blood was determined by monitoring for cells expressing the CD45.1 allele (donor) and CD45.2 allele (host) in B-cells, T-cells and granulocytes. Donor and host granulocyte chimerism is illustrated.

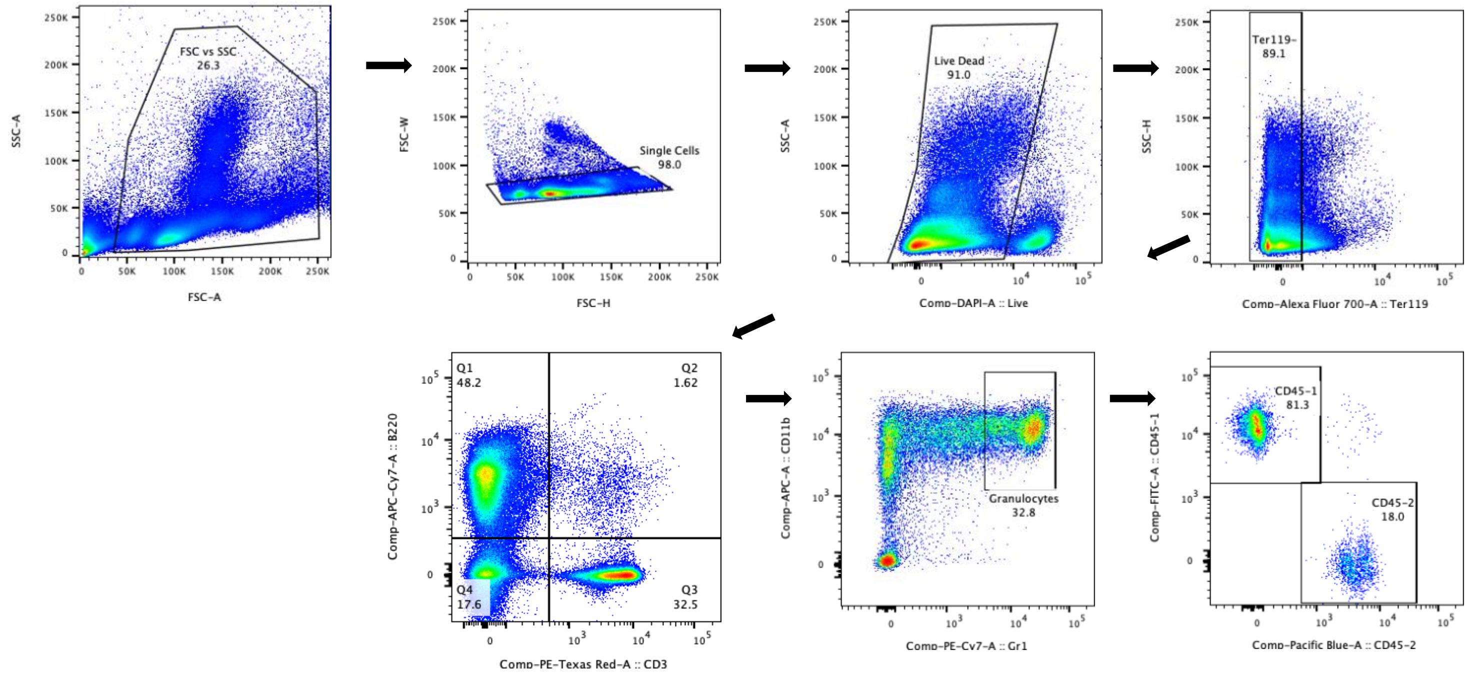

**S3: Representative gating strategy for long-term (LT) hematopoietic stem cells (HSC) chimerism in the bone marrow.** Post-IUHSCT, donor engraftment in the bone marrow was determined by monitoring for cells expressing the CD45.1 allele (donor) and CD45.2 allele (host) in Lin-CD117+Sca1+CD150+CD48- LT-HSC.

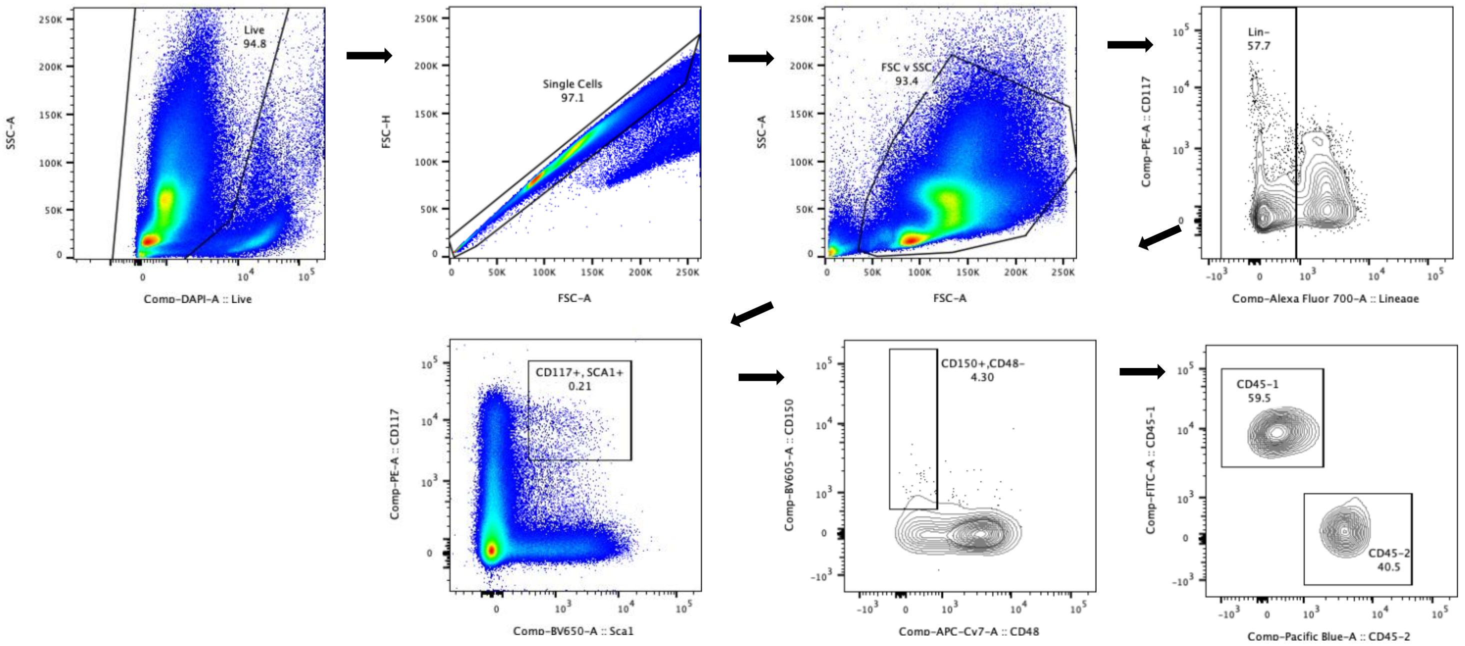


**Supplemental Materials:**

| | | |
|---|---|---|
| **Pacific Blue™ anti-mouse CD45.1 Antibody** | **A20** | **BioLegend** |
| **FITC anti-mouse CD45.2 Antibody** | **104** | **BioLegend** |
| **Alexa Fluor® 700 anti-mouse TER-119/Erythroid Cells Antibody** | **TER-119** | **BioLegend** |
| **PE/Dazzle™ 594 anti-mouse CD3 Antibody** | **17A2** | **BioLegend** |
| **Brilliant Violet 785™ anti-mouse CD19 Antibody** | **6D5** | **BioLegend** |
| **Brilliant Violet 605™ anti-mouse CD8a Antibody** | **53-6.7** | **BioLegend** |
| **PE anti-mouse CD4 Antibody** | **GK1.5** | **BioLegend** |
| **APC/Cyanine7 anti-mouse/human CD45R/B220 Antibody** | **RA3-6B2** | **BioLegend** |
| **APC anti-mouse/human CD11b Antibody** | **M1/70** | **BioLegend** |
| **APC/Cyanine7 anti-mouse Ly-6G/Ly-6C (Gr-1) Antibody** | **RB6-8C5** | **BioLegend** |
| **PE anti-mouse CD117 (c-kit) Antibody** | **2B8** | **BioLegend** |
| **Alexa Fluor® 700 anti-mouse Lineage Cocktail with Isotype Ctrl** | **17A2; RB6-8C5; RA3-6B2; Ter-119; M1/70;** | **BioLegend** |
| **Brilliant Violet 650™ anti-mouse Ly-6A/E (Sca-1) Antibody** | **D7** | **BioLegend** |
| **APC/Cyanine7 anti-mouse CD48 Antibody** | **HM48-1** | **BioLegend** |
| **Brilliant Violet 605™ anti-mouse CD150 (SLAM) Antibody** | **TC15-12F12.2** | **BioLegend** |
| **LIVE/DEAD™ Fixable Blue Dead Cell Stain Kit, for UV excitation** | | **ThermoFisher** |
